# Assignment of the resting state of the secondary active transporter BetP by integrating spectroscopic measurements and molecular simulations

**DOI:** 10.1101/307710

**Authors:** Vanessa Leone, Izabela Waclawska, Katharina Kossmann, Caroline Koshy, Monika Sharma, Thomas F. Prisner, Christine Ziegler, Burkhard Endeward, Lucy R. Forrest

## Abstract

The glycine betaine symporter BetP regulates the osmotic stress response of *Corynebacterium glutamicum*, a soil bacterium used extensively in biotechnology. Although BetP is a homotrimer, biochemical studies have shown that each protomer is able to transport its substrate independently. Crystallographic structures of BetP have been determined in several conformations, seemingly capturing outward-open, inward-open and occluded states, both loaded with the substrate and in the apo form. However, it has been challenging to establish a correspondence between each of these structures and specific states in the mechanism of the transporter under more physiological conditions. To this end, we examined the dynamics of spin-labelled BetP using pulsed electron-electron double resonance (PELDOR) under different stimuli. We then carried out molecular simulations of structures of the BetP monomer to interpret the PELDOR data, using the enhanced-sampling methodology EBMetaD (1), whereby the dynamics of the protein are minimally biased so as to reproduce the experimental data. Comparison of the magnitude of the biasing work required for different input structures permitted us to assign them to specific states of the transport cycle under each of the experimental conditions. In particular, this analysis showed that BetP adopts inward-facing conformations in the presence of excess sodium, and a mixture of states when betaine is added. These studies better delineate the major conformations adopted by BetP in its transport cycle, and therefore provide important insights into its mechanism. More broadly, we illustrate how integrative simulations can aid interpretation of ambiguous structural and spectroscopic data on membrane proteins.

Abbreviations

BetP: betaine permease
TM: transmembrane
POPG: palmitoyl oleyl phosphatidyl-glycerol
DDM: β-dodecyl-maltoside
SEC: size-exclusion chromatography
EPR: electron paramagnetic resonance
PELDOR: pulsed electron-electron double resonance
DEER: double electron-electron resonance
EBmetaD: ensemble-biased metadynamics
MD: molecular dynamics
R5: 1-oxyl-2,2,5,5-tetramethylpyrrolidin-3-yl)methyl methanethiosulfonate

## Introduction

Secondary active transporters catalyze the accumulation of substrates across biological membranes, often coupling this process to the energetically downhill transport of ions. To do so, these transporters cycle through a series of conformations whereby the binding sites for the translocated species are alternatively exposed to each side of the membrane, in a mechanism known as ‘alternating access’ (2).

One of the best studied superfamilies of secondary-active transporters adopts what is known as the LeuT fold. This protein architecture consists of two repeats of five transmembrane helices, where the two repeats are inserted in the membrane in opposite orientations (3). Proteins with this architecture include sodium-coupled transporters, such as LeuT (4), from the neurotransmitter:sodium symporter (NSS) family; Mhp1 (5) of the nucleobase:cation symporter-1 (NCS1) family; and BetP (6) from the betaine/choline/carnitine transporter family (BCCT). While these proteins share a common fold and coupling ion, the transport stoichiometry varies; substrate transport by Mhp1 (5) is coupled to the translocation of one sodium ion, which binds at the so-called Na2 site, while LeuT and BetP both couple transport to the flux of two sodium ions, with the additional ion binding to the Na1 or Na1’ site, respectively (4, 7–9).

Unlike other transporters with the LeuT fold, BetP forms constitutive trimers. Although each BetP protomer is a functional transporting unit, trimerization is required for the response to osmotic stress, which is often referred to as activation (10). This activation process involves the N- and C-terminal domains of each protomer (11) and modulates (accelerates) the transport kinetics in a manner that depends on intracellular potassium concentration (12) and lipid composition (13). However, activation is not required for the Na+-coupled uptake of betaine, which occurs at a basal level in the absence of potassium or osmotic stimulation.

Through X-ray crystallography, BetP has been captured in almost all expected conformations along the transport cycle (6, 14–16), indicating conformational changes between transmembrane helices (TM)^1^ 1’, 5’, 6’ and 10’ (16), relative to the helices in the so-called hash domain (TM 3’-4’ and 8’-9’, Figure 1). In spite of the unprecedented insights into the conformational cycle of BetP garnered from these structures, the question remains how the interconversion between states is controlled by ion and substrate binding. Spectroscopic methods have great potential to fill that gap. For example, pulsed electron-electron double resonance (PELDOR) spectroscopy, also known as double electron-electron resonance (DEER), has provided important insights into how coupled ions and substrate influence the conformational preferences of LeuT and Mhp1 (17, 18). Specifically, Mchaourab and collaborators have shown that while the presence of Na+ is able to shift LeuT towards outward-facing conformations, Mhp1 requires also substrate binding to stabilize outward-facing conformations albeit while sampling a small population of inward-facing conformations. The different effect of sodium ions on selecting transporter conformations, together with the knowledge that only LeuT has an Na1 binding site, led to the hypothesis that Na1 (and not Na2) binding is required for the inward-to-outward conformational change (17, 18). However, biochemical accessibility studies of LeuT upon modification of the two binding sites indicated that Na2 suffices to control the inward-to-outward transition (19), suggesting that other properties of Mhp1 instead contribute to its unexpected conformational preferences.

**Figure 1.**
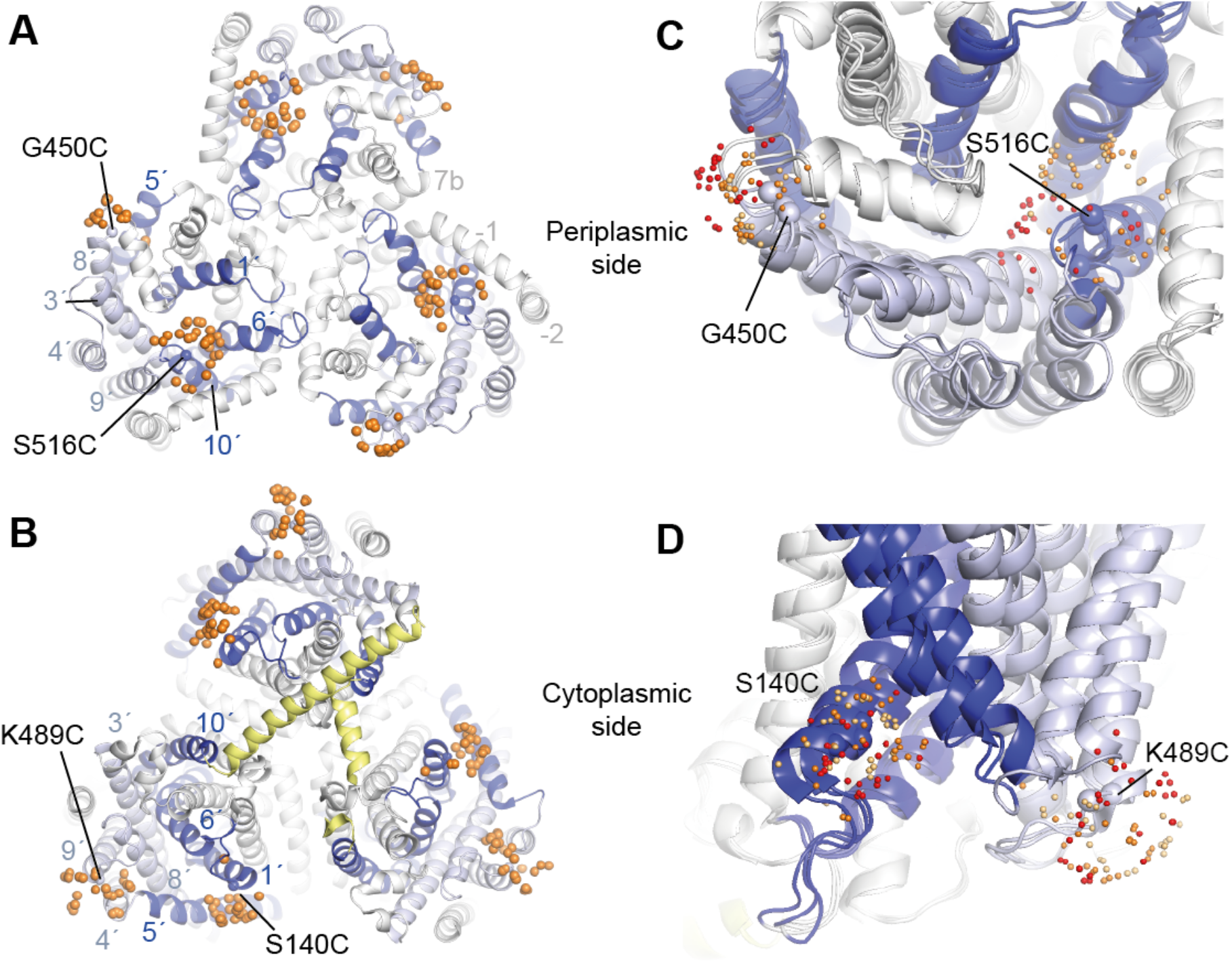
Structure of BetP and spin label positions. The structure of BetP shown as cartoon helices, highlighting transmembrane helices 1’, 5’, 6’ and 8’ lining the substrate pathway (*blue*), helices 3’, 4’, 8’ and 9’ in the so-called hash domain (*pale blue*), and the cytoplasmic C-terminal extension (*yellow*). Helices −2, −1 and 7b (*white*) are also labelled. (**A, B**) Trimer structure of BetP (PDB code 4C7R). Each BetP protein chain was labelled at two cysteines on the periplasmic surface, introduced at positions G450 an S516 (A) or on the cytoplasmic surface at positions S140 and K489 (B). Ca atoms of labelled positions are shown as spheres. Position of the nitroxide oxygen atom in the MTSSL spin label in orientations predicted using a rotamer library is shown as small orange spheres. (**C, D**) Comparison of the predicted spin-label positions in structures of the three major conformations of BetP: outward-open (PDB code 4LLH chain A); closed (PDB code 4AIN chain A); or inward-open (PDB code 4C7R chain A). Nitroxide N atom positions are colored red, orange and yellow, respectively, for the three conformations.

In PELDOR, a spectroscopic signal resulting from the interaction between two spin labels in a molecular ensemble is used to derive a probability distribution of the spin-to-spin distance. Typically, a pair of nitroxide radicals are covalently attached to cysteine residues introduced at specific positions in the protein using site-directed spin labeling (SDSL). This kind of spin label is relatively flexible, however, posing a challenge for the interpretation of these distance distributions in terms of specific protein conformations. In principle, molecular dynamics simulations provide a means to overcome this challenge, as they can be used to generate ensembles of spin-label configurations for different protein conformations. Unfortunately, however, conventional molecular dynamics simulations will typically fail to reproduce the probability distributions derived from experiment, either due to insufficient computational sampling, i.e., limited simulation time, or because the starting structure used in the simulation is not a major contributor to the experimental data. Thus, advanced simulation strategies have been developed to ensure that such simulations reproduce the experimental distribution directly, with the minimum perturbation relative to conventional simulations (20). However, the high computational cost of these strategies makes full atomistic detail impractical (17, 18). More recently, a variation of this maximum-entropy approach, known as EBMetaD was developed to make this kind of calculation affordable in atomistic detail (1).

Thus, to examine how the conformational landscape of BetP is influenced by ion and substrate binding, we performed PELDOR measurements for SDSL BetP proteoliposomes, in the presence of different ion and substrate concentrations, and analyzed the experimental results with EBMetaD simulations of different X-ray structures of the protein. This strategy allowed us to rank each simulated structure based on the ease or difficulty of targeting the experimental distribution. In this way, we examined which of these structures is most representative of the experimental conditions associated with each of the PELDOR measurements.

By the use of this integrative approach, we show that BetP is thermodynamically controlled in a similar manner as Mhp1, but differently from LeuT. Specifically, the presence of the substrate plus sodium ions, appears to foster a mixture of inward- and outward-facing states of BetP, whereas in the absence of betaine, sodium favors an inward-facing state. We discuss the implications of these finding in the context of the osmotic stress-dependent activation mechanism of BetP.

## Materials and Methods

### Spin label rotamer prediction

The range of possible spin-spin distance values was estimated using the multiscale modeling of macromolecular systems (MMM2015.1) software package (21–23) to model all possible rotamers of R1 probes attached to either cytoplasmic (S140 and K489) or periplasmic (G450 and S516) sides of BetP. To assess the range of distances arising from spin coupling across protomers, we predicted the rotamers of the spin label on the inward-open trimer (PDB entry 4C7R (14)). To predict distance distribution changes upon conformational change within a protomer, spin label rotamers were modelled onto structures of the outward-open (chain A of PDB entry 4LLH (15)), closed (chain A of PDB entry 4AIN (16)), or inward-open (chain A of PDB entry 4C7R (14)) states. Since the standard spin label rotamer libraries (R1A_175K and R1A_298K) included, in some cases, only a few rotamers, we used the R1A_298K_xray library (23). Note that the scope of these calculations is not to predict the exact distance distribution of each conformation, but rather the range of possible distances, so as to facilitate separation of the entire experimental distance distribution into intra- and inter-protomeric spin-spin distances.

### Site-directed mutagenesis

Site-directed mutagenesis was performed with the QuickChange Site-directed Mutagenesis Kit II (Stratagen) and PfuUltra DNA polymerase in pASK-IBA5betP plasmid (24). All the plasmids were fully sequenced to confirm the specific mutation.

### Protein expression and purification

Wild type, cysless, and cysteine variants of BetP were produced and purified as described previously (25) using the primers given in Table S1. Briefly, pASK-IBA5betP wild type and mutants were transformed and heterologously expressed in *E. coli* One Shot^®^Invitogen DH5α™-T1. The cells were grown in LB medium with carbenicillin (50 μg/ml) at 37 °C. Protein expression was induced with anhydrotetracycline (200 μg/l). After centrifugation at 4°C, the cells were resuspended in a buffer containing 50 mM Tris-HCl, pH 7.5, 17.2% glycerol, and 1 mM protease inhibitor pefabloc. Membranes were isolated from broken cells by centrifugation and, subsequently solubilized with 1% β-dodecyl-maltoside (DDM). Solubilized membranes were loaded on a pre-equilibrated Strep-Tactin Macroprep (IBA) column and washed with 40 column volumes of 50 mM Tris-HCl, pH 7.5, 200 mM NaCl, 8.6% glycerol, and 0.1% DDM. The protein was eluted in the same buffer supplemented with 5 mM desthiobiotin. All purification steps were performed at 4°C. BetP was further purified to remove the unbound spin label by size-exclusion chromatography (SEC). The protein solution was loaded in a superose 6 10/300 column connected to an Äkta system; the column was pre-equilibrated with 25 mM Tris-HCl, pH 7.5, 200 mM NaCl and 0.1% DDM.

### Uptake assays

The protein was reconstituted in *E. coli* polar lipid extract (Avanti) as described (25, 26). Liposomes (20mg/ml) were prepared by extrusion through a filter (polycarbonate membrane, pore size of 400nm, Avestin) and diluted 1:4 in 100 mM KP_i_ pH 7.5. The solution was titrated with 10% (w/v) Triton X-100 and then mixed with the purified protein at a lipid to protein ratio of 30:1 for uptake experiments. The detergent was removed by adding BioBeads SM-2 (Bio-Rad) at ratios (w/w) of 5 (BioBeads/Triton X-100) and 10 (BioBeads/DDM) in 5 steps. The proteoliposomes were centrifuged, washed, and resuspended in 100 mM KP_i_ pH 7.5 buffer to a concentration of 60 mg/ml before freezing in liquid nitrogen, and storing at −80 °C.

Uptake of [^14^C]-labeled glycine betaine were performed as described (25). Briefly, proteoliposomes were extruded (pore size 400 nm, polycarbonate membrane, Avestin) and centrifuged. Afterward they were resuspended in internal buffer (100 mM KP_i_ pH7.5) to a lipid concentration of 60 mg/ml. Uptake measurement was initiated by diluting proteoliposomes at a ratio of 1:200 in the external buffer (20 mM NaP_i_ pH 7.5, 25 mM NaCl, 50 μM [^14^C]-labeled glycine betaine, and 1 μM valinomycin). The external osmolarities were adjusted to 0.6 osmol/kg with proline. Samples were filtered for various times through nitrocellulose filters, and the amount of [^14^C]-glycine betaine incorporated into the proteoliposomes during uptake was determined by scintillation counting.

### Site directed spin labeling

Cysteine variants of BetP were labeled with a 30-fold molar excess of the spin label (1-oxyl-2,2,5,5-tetramethylpyrrolidin-3-yl)methyl methanethiosulfonate, R5). Free spin label was removed by SEC as described in the Protein Expression and Purification section. The spin label concentration was estimated by continuous wave (cw) EPR measurements performed at X-band frequency (9.4 GHz) and used to estimate labeling efficiency based on the protein concentration determined using amido black. Free spin labels were observed at <5% in the sample after SEC purification.

### Protein reconstitution for PELDOR measurements

Similar to the protein reconstitution for uptake, the spin labeled protein was added to extruded liposomes (in 200 mM Tris-HCl, pH 7.5) with a lipid to protein ratio of 20:1. After incubation, the lipid/protein mixture was transferred to a dialysis membrane (MW cutoff 12-14 kDa, Spectrum labs). BioBead SM-2 at ratios (w/w) of 5 (BioBeads/Triton X-100) and 10 (BioBeads/DDM) was added to the dialysis buffer in 4 steps. The proteoliposomes were centrifuged and resuspended in two different buffers, for the experimental conditions assessed, prepared in deuterated water; 200 mM Tris-HCl pH 7.5, 500 mM NaCl or 200 mM Tris-HCl pH 7.5, 300 mM NaCl, 5 mM betaine. After an additional extrusion and centrifugation step, the proteoliposomes were again resuspended in the corresponding buffers.

### PELDOR measurements

EPR measurements were performed under two experimental conditions, either a saturating concentration of sodium (500 mM NaCl) or saturating concentrations of both sodium and betaine (300 mM NaCl, 5 mM betaine). Measurements were performed with two spin label pairs; S140R5/K489R5 and G450R5/S516R5, both in the background of the C252T cys-less variant.

All PELDOR data were recorded on an ELEXSYS E580 EPR spectrometer (Bruker) equipped with a PELDOR unit (E580–400U, Bruker), a Bruker-D2 resonator for Q-Band frequencies using a 10W Amplifier, a continuous-flow helium cryostat (CF935, Oxford Instruments), and a temperature control system (ITC 502, Oxford Instruments). A dead-time free, four-pulse sequence was used with phase-cycled π/2, ESSEM averaging (8×16ns), 20ns pump, and 32 ns detection pulses (27).

### PELDOR analysis

The use of a 6-spin system raises the possibility of ghost signals due to multi-spin effects, which might mask peaks or create additional peaks at short distances (28, 29). We therefore took steps during data processing to mitigate such effects (see **Supporting Information**). In addition, rotamer library-based modeling indicates that these effects will be negligible in our case (see **Supporting Information, Figure S1**).

### Molecular dynamics simulation setup

Simulations were performed on monomers isolated from three different trimeric crystal structures, representing the substrate-free inward conformation (PDB entry 4C7R chain C, 2.7 Å resolution (14)), the substrate bound inward-open state (PDB entry 4AIN chain C, 3.1 Å resolution (16)), and outward-open substrate-free and substrate-bound states (PDB entry 4LLH chain A, 2.95 Å resolution (15)). To reproduce the experimental conditions, we modified the ion/substrate occupancy accordingly; two sodium ions were added to the crystal structure of the apo inward-facing (4C7R), betaine-bound inward-facing (4AIN), and choline-bound outward-facing (4LLH) conformations (see below). The outward-open crystal structure bears a point mutation of residue Gly153 to an aspartate, which was mutated back to the wild type sequence (D153G). For the substrate-bound simulation, a betaine molecule was positioned at the choline binding site by superposing the trimethylamine group. Glu161 and Asp239 were assumed to be neutral, based on a combination of pKa calculations performed on the X-ray structures using MCCE (30) (**Table S2**) and analysis of the stability of contacts during molecular simulations (not shown). However, for Asp470, the predicted *pK_a_* of Asp470 was close to 7 for some states (**Table S2**), and therefore for every conformation, we generated two systems: one with Asp470 protonated and the other with Asp470 deprotonated.

All proteins were embedded in a hydrated (~15,000 waters) bilayer of 219 palmitoyl oleyl phosphatidyl-glycerol (POPG) lipids with GRIFFIN(31). All systems were set to neutral by modifying the number of sodium (254-259) and chloride ions (25–30) in the bulk water.

Each system was subsequently equilibrated for 20 ns by a series of restrained simulations, involving 2 ns trajectories with stepwise release of harmonic positional restraints, as follows: a) 15 kcal/mol A^−2^ applied to side chains, backbone, and ligands; b) backbone and ligand restraints of 15 kcal/mol A^−2^ and side chain restraints of 4 kcal/mol A^−2^; c) Ca atoms and ligand restraints of 15 kcal/mol A^−2^, backbone restraints of 4 kcal/mol A^−2^ and side chain restraints of 1 kcal/mol A^−2^; d) ligand restraints of 15 kcal/mol A^−2^, Ca atom restraints of 4 kcal/mol A^−2^, and side chain restraints of 1 kcal/mol A^−2^. In the final 4-ns long step, the Ca atoms restraint was 1 kcal/mol A^−2^, and the ligand restraint was 4 kcal/mol A^−2^. This was followed by 108 ns of unconstrained equilibration. Pairs of R5 spin labels were then added at Ser140 and Lys489 (cytoplasmic pair) or at Gly450 and Ser516 (periplasmic pair) using the CHARMM-GUI (32). The labeled system was further equilibrated for 100 ns.

In all simulations, to prevent substrate and ion unbinding from the simulated open structures, while not explicitly defining their exact (unknown) coordination, we loosely restrain each substrate to its cavity by applying restraints based on a coordination number (*C_num_*) variable defined as:

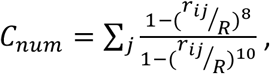

where the *i* index refers to the ligand (sodium or betaine nitrogen atom) and *j* indicates the protein atoms coordinated to this ligand. The Na1 site was defined as the side-chain oxygen of residues Thr250 and Thr246 and the backbone oxygen of Thr246 (8). The Na2 site was defined as the oxygen of the side chains of residue Thr467 and Ser468 and the backbone side chain of residue Met150. The betaine nitrogen atom was coordinated with the Cγ atom of Trp377, the Oη atom of Tyr197 and the backbone oxygen of Ala148. The cutoff, *R*, was set to 2.5 for sodium coordination and to 2.0 for betaine coordination. In addition, a harmonic restraint of 50 kcal/mol was applied that prevents the Na1 coordination from adopting values <1.3 Å, Na2 coordination <2.1 Å and betaine coordination <2.0 Å.

Molecular dynamics simulations were performed using NAMD v2.9 (33). The Ensemble-Biased Metadynamics (EBMetaD) was performed with a modified version of PLUMED v1.3 (34) provided by Fabrizio Marinelli and José Faraldo-Gómez. The NAMD collective variables module (35) was used to define the ligand coordination number variables.

The CHARMM36 force field (36–38) was used for the protein, lipid and ions; TIP3P for the water molecules (39); and betaine parameters from Ma et al (40). The simulations were performed at constant temperature (298 K) and pressure (1 atm), imposed with a Nosé-Hoover Langevin barostat and thermostat. The membrane area was kept constant and periodic boundary conditions were used in all directions. The time step of the simulations was 2 fs. Electrostatic interactions were calculated using particle mesh Ewald with a real-space cutoff of 12 Å. The cutoff of van der Waals interactions was also set to 12 Å. A switching function starting at 10 Å was applied to both electrostatic and van der Waals interactions.

### Biasing the simulated ensemble to the experimental distance distribution

The target distance distribution was enforced using EBMetaD (1) on the centers of mass of the two spin label nitroxide bonds. Briefly, this method adds an adaptive biasing potential in the form of additive Gaussians to the underlying force field during the simulations. Unlike canonical metadynamics, the height of each applied Gaussian depends on the probability of the targeted distribution, such that the Gaussian will be larger for low probability regions and smaller for high probability regions. Importantly, the bias introduced during the simulations is the minimum necessary to fulfill the target distribution, preventing overfitting to the distance data.

### Analysis of the applied work and biased trajectories

From the bias potential applied during a simulation we define the work along the distance distribution, *W* to be:

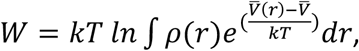

where *r* is the nitroxide-nitroxide distance, *ρ*(*r*) is the experimental distance distribution being targeted, *k* is the Boltzmann constant, and *T* is the temperature. The quantity 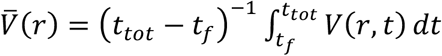 is the average bias potential added during a simulation of length *t_tot_*, in which *V*(*r, t*) is the bias potential added at time *t*. Here, 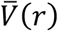 is averaged over the last 0.8 μs of each 1-μs enhanced simulation, i.e., excluding the initial “filling” time, *t_f_* = 0.2 μs. The offset term 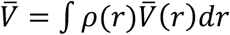 is defined as the mean value of 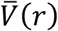 over the experimental distance distribution and allows for comparison between simulations with different starting structures.

To evaluate the effect of the bias on the distance distributions, de-biased distance distributions were computed as follows:

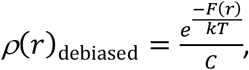

where 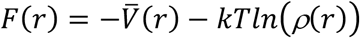 is the free energy of an unbiased simulation as a function of the spin-label distance, *r* (eq. 6 in (1)), and *C* is a normalization factor, 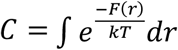.

To isolate distances potentially arising from inter-protomer coupling, we computed the work required to bias the trajectory to the distribution only in a specific distance range. To this end, in the work calculation we considered a modified average bias potential, 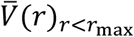, that was assumed to be flat for distances *r* above a threshold distance, *r*_max_, which was set to 37 Å. In this case, we also modified the reference experimental distribution (*ρ*_modified_) to account for the bias potential change. Thus,

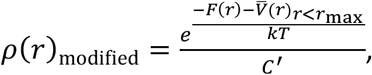

where *C*′ is a normalization factor, 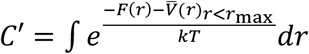.

## Results

### BetP retains functionality upon introduction of periplasmic or cytoplasmic spin-labels

Nitroxide radicals were introduced at different positions of the cysteine-less BetP mutant C252T (11, 25) by site-directed spin labeling. BetP is a homotrimer (**Figure 1**), but each of the protomers is functional, and operates independently (41). Therefore, two spin labels must be introduced on each protomer to report on transport-related conformational changes. Here, we covalently linked methyl methanethiosulfonate R5 spin labels to two different residues in each protomer, resulting in a total of six spins. We designed these systems based on three considerations. First, the spin-labels should report on distances between helices involved in the conformational changes implied by available crystal structures (**Figure 1C-D**), namely TM helices 1’, 5’, 6’ and 10’ (16), relative to TM helices 3’-4’ and 8’-9’ (i.e., the so-called hash domain). Second, the intra-protomer spin-spin distances should be shorter than the inter-protomer distances, to facilitate the interpretation of the results. Finally, the labelled protein should be functional.

Comparison of the known structures of BetP suggests that the transport cycle involves structural states that differ only by ~ 5 Å. This relatively small magnitude, combined with the three requirements mentioned above, limited the number of possible positions for the spin labels. Among the positions tested (**Table S3**), we ultimately used S140R5/K489R5 to track possible conformational changes of TM 1’ relative to TM 9’, which opens the cytoplasmic pathway in the crystal structures (**Figure 1B, 1D**). On the periplasmic surface of the transporter, we used G450R5/S516R5 to assess the displacement of TM 10’, which gates the periplasmic pathway relative to TM 8’ (**Figure 1A, 1C**). For both pairs, R5 labelling was achieved with high efficiency (**Table S3**). [^14^C]-betaine uptake measurements of spin-labelled proteoliposomes show both constructs are functional: S140R5/K489R5 retains activity but at a diminished rate, whereas the activity of G450R5/S516R5 is only slightly decreased compared to the wildtype (**Figure 2**).

**Figure 2:**
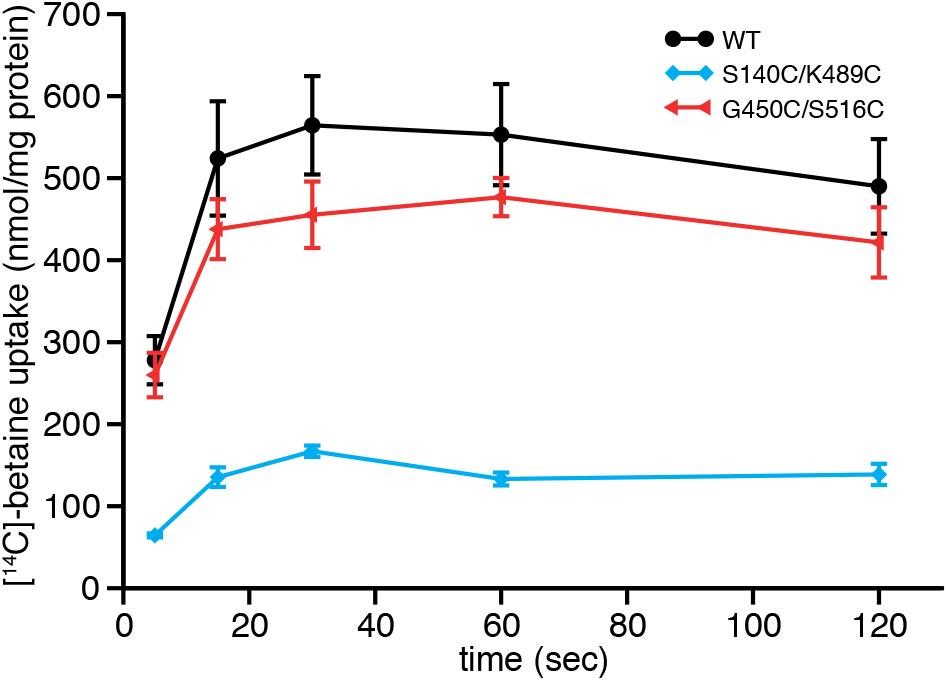
Betaine uptake of spin-labeled BetP variants in *E. coli* polar lipid proteoliposomes. Uptake of betaine in nmol per mg protein was measured at 0.6 osmol/kg as a function of time for R5-labeled BetP cysteine variants reconstituted into *E. coli* polar lipid liposomes. Uptake was initiated by adding saturating concentrations of 50 μM [^14^C]-betaine. Each value is the mean ± S.E.M. of at least six independent measurements.

### Distance between periplasmic spin labels differs in presence or absence of betaine

Having confirmed the functionality of our two spin-labeled constructs, we then measured the interaction between the spins using PELDOR (**Figure 3**). The measurements were performed in proteoliposomes (42), which represent a more native environment than, say, detergent micelles. We specifically asked how the conformational preferences of the transporter are influenced by the presence of its two substrates, sodium and betaine. We therefore tested two reference conditions, namely a saturating concentration of sodium (500 mM NaCl inside and outside the proteoliposomes) and saturating concentrations of both sodium and substrate (300 mM NaCl, 5 mM betaine, inside and outside the proteoliposomes). For the cytoplasmic spin-label pair (S140R5/K489R5), the distance distributions are very broad, and are comparable in the two conditions tested (**Figure 3B**). This result suggests either that the conformation of the protein in this region does not change measurably under these two conditions, or that this conformational change is masked by the intrinsic flexibility of the spin labels. By contrast, the distribution of distances for the periplasmic pair (G450R5/S516R5) features an additional peak at shorter distances (r < 30 Å) when both betaine and sodium are present compared to sodium alone (**Figure 3D**). A relative displacement of G450 and S516 is consistent with crystal structures of BetP in inward- and outward-facing states; however, the distance change between the respective Cα atoms that is implied by these structures is only ~2 Å (PDB 4C7R_A at 28.7 Å and PDB 4DOJ_B at 30.6 Å, respectively), i.e. a value much smaller than the overall width of the PELDOR distance distributions (**Figure 3D**). Based on this data, it is therefore non-trivial to establish a correspondence between a given substrate condition and the preferred structural state of the transporter.

**Figure 3.**
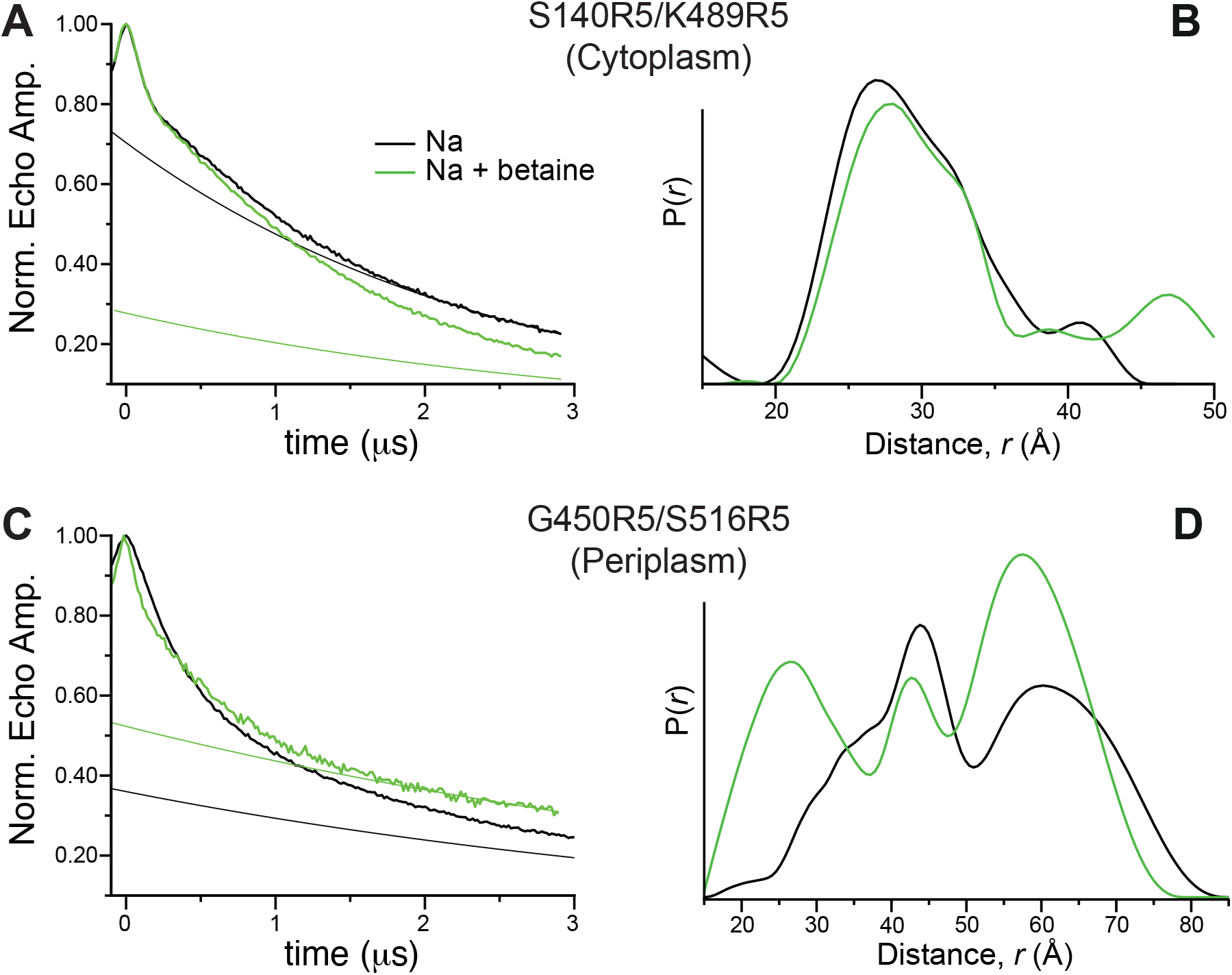
PELDOR data for spin label pairs introduced in the BetP trimer. **(A, C)** PELDOR time traces and assumed background traces (thinner smooth lines) measured in the presence of 500 mM NaCl (*black*) or 300 mM NaCl and 5 mM betaine (*green*), for a pair of spin labels introduced (**A**) on the cytoplasmic surface between TM 1’ and TM 9’ (S140R5 and K489R5) or (**C**) on the periplasmic surface between TM 8’ and TM 10’ (G450R5 and S516R5). **(B, D)** Probability of a distance *P*(*r*) versus distance (*r*) between spins on the cytoplasmic (B) or periplasmic (D) sides of BetP. Distance distributions were derived from Tikonov regularization of the PELDOR time traces in A and C.

### EBMetaD simulations of the BetP protomer converge on the experimental distribution

To formulate a structural interpretation of the PELDOR data, we resorted to using molecular dynamics simulations. Specifically, we employed a recently developed enhanced-sampling technique known as EBMetaD (1) to construct conformational ensembles that reproduce the PELDOR distance distributions, using different crystal structures as the starting point. We then asked which of these input structures is most representative of the calculated ensemble representing the experiment.

BetP forms homotrimers, and the trimerization interface is believed to regulate the kinetics of the transporter in a K^+^-dependent manner. However, the monomeric protein is stable and fully functional. Therefore, for simplicity and maximal computational efficiency, the simulation system used here is a BetP monomer and not a trimer. Since the PELDOR measurements were performed on the trimer, we modified the experimental distance distributions by excluding contributions within the range of values that could only correspond to interactions between spin labels in different protomers. To identify this distance range, we modeled and analyzed a diverse ensemble of spin-label rotamers at either cytoplasmic (S140 and K489) or periplasmic (G450 and S516) positions (see Methods) and computed the distances between the nitroxide groups in spin-label pairs in the same protomer, for the outward-facing, inward-facing, and closed conformations of the transporter (**Figure S2A, S2C**). Based on this analysis, we concluded that the maximum possible distance for intra-protomer spin coupling is ~45 Å. Thus, the peak at ~60 Å observed in the PELDOR distance distributions (**Figure 3D**) likely reflects the interaction between labels in different protomers. We therefore filtered out this peak by fitting a Gaussian curve to it and subtracting that Gaussian from the experimental data (**Figure S2E, S2F**). Although the result of this procedure is only an approximation, it is worth noting that the region of the distribution that is altered (peak ~42 Å) is similar in the two experimental conditions considered, which is therefore not very informative. The most important differences are observed in the 15-30 Å range (**Figure 3D**), which is not influenced by this filtering procedure. Thus, comparison of simulations guided by the filtered distributions is a valid strategy to establish the desired correspondence between conformational states and specific experimental conditions.

Using the filtered distance distributions as target, EBmetaD simulations were initiated for structures of either the outward- or the inward-facing state, and either with two sodium ions bound or with two sodium ions plus a betaine molecule bound, depending on the experimental condition being compared. Finally, since the protonation state of Asp470 was found to be sensitive to the structure of the protein (**Figure 4, Supporting Information**), each simulation was repeated with Asp470 either protonated or deprotonated.

**Figure 4.**
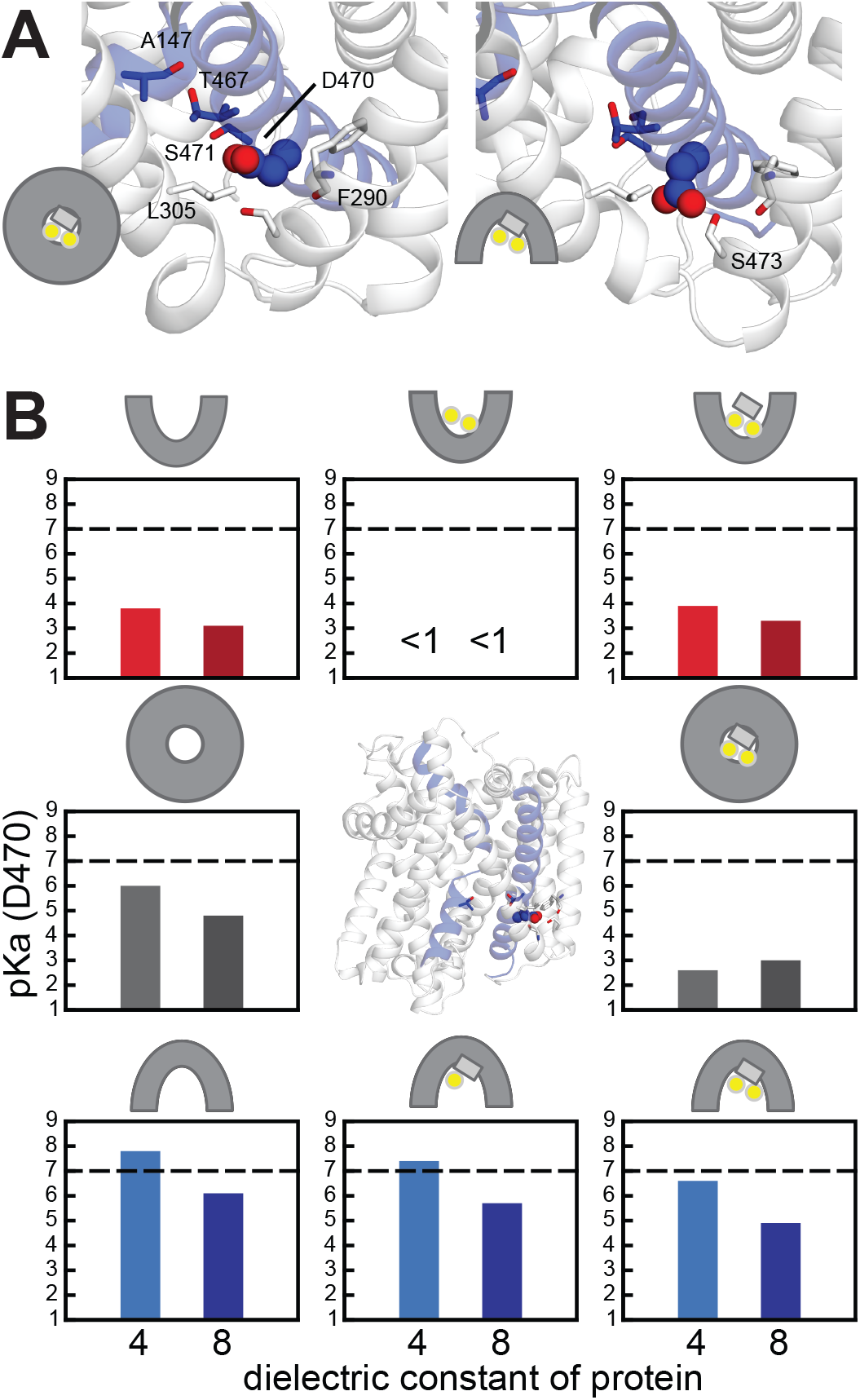
The protonation state of Asp470 of BetP may vary during the transport cycle. **(A)** Structures of BetP showing the two major rotamers observed for Asp470 (*spheres*), facing toward Ser293 in the TM4’-5’ loop in the inward-facing structures, or toward Ser471 in TM8’ in the closed (4AIN_B) or inward-facing (4AIN_C) structures. Key residues are shown as sticks. Ala147 and Thr467 form the Na2 binding site. **(B)** pKa values were computed for different structures in the physiological transport cycle, with the continuum electrostatics method MCCE (30) and with the protein dielectric constant set to 4 or 8. Top row: The outward-open structures used were 4DOJ_B for the apo state; 4DOJ_B with sodium added and energy minimized; and 4LLH_B for the fully-loaded state. Middle row: the closed structures used were 4AIN_A for the apo state and 4AIN_B for the holo state. Bottom row: inward-open structures used were 4C7R_C for the apo state, 4AIN_C both for the fully-loaded state, and the betaine plus Na1’ state. Data for the outward-facing conformations are taken from (15), and were obtained from the crystal structure of G153D mutant that was *in silico* mutated back to the wildtype sequence, while the choline molecule was substituted by a betaine molecule. The central figure indicates the position of D470 (*spheres*) close to the cytoplasmic pathway in the inward-open structure 4C7R_A. TM 1’ and TM 8’ are colored blue.

To assess whether these simulations were sufficiently long, we compared the distribution of spin-label distances obtained during the simulations with the experimental distance distribution. After 1 μs of sampling time, all the simulations had converged to match exactly the experimental distance distributions, even for the more challenging case in which the distribution includes two peaks (**Figure 5**).

**Figure 5.**
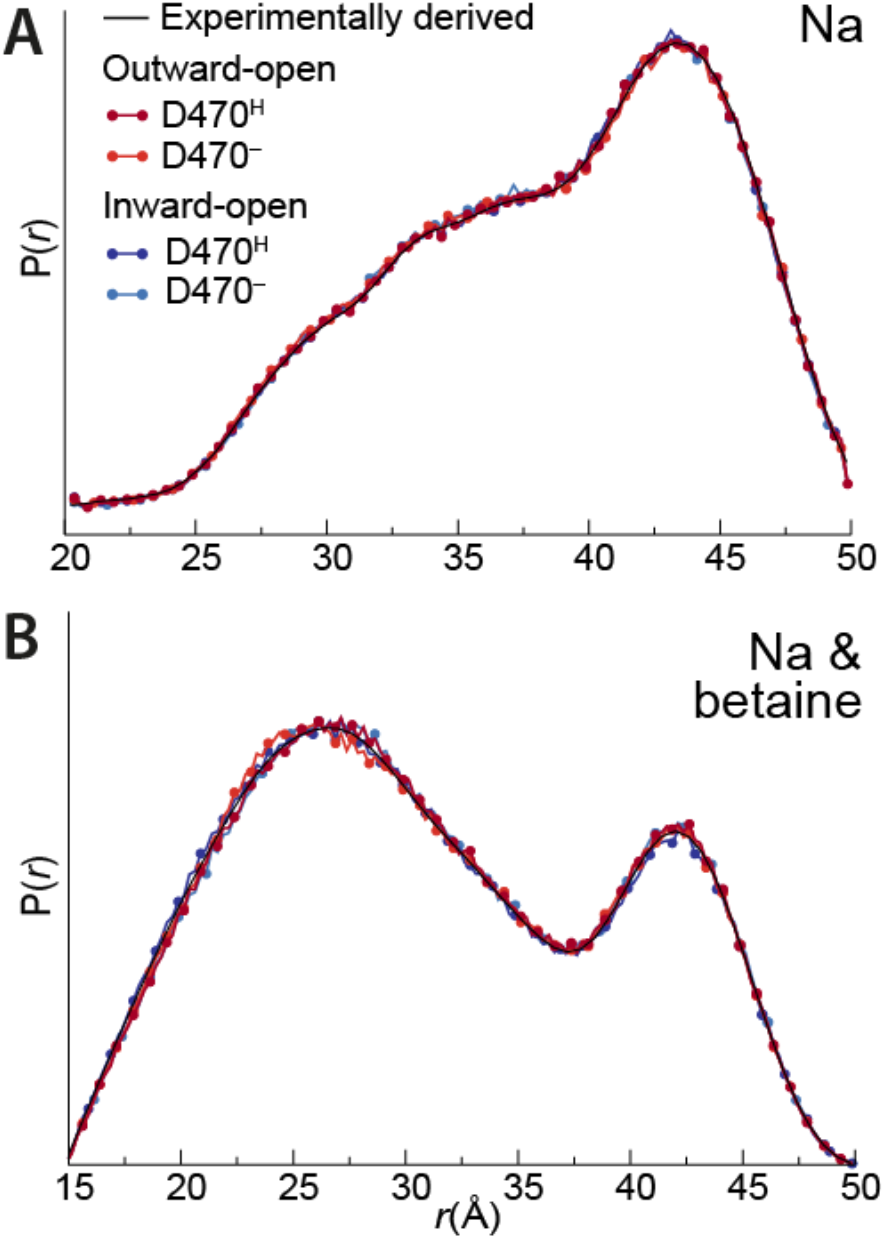
Convergence of simulated to experimental distance distributions for the periplasmic spin label pair. The probability of a distance *P*(*r*) is plotted versus distance (*r*). The PELDOR-based distances (*black lines*), measured in the presence of 500 mM NaCl (**A**) or 300 mM NaCl plus 5 mM betaine (**B**), are compared with distances obtained in 1 μs-long EBMetaD molecular dynamics simulations, performed for BetP monomers in the presence of two sodium ions (**A**) or with two sodium ions plus a betaine substrate (**B**). The simulation was started with BetP structures of either inward-facing (PDB code: 4C7R, *blue*), or outward-facing conformations (PDB code: 4LLH chain A, *red*), with Asp470 either protonated (*darker colors*), or deprotonated (*paler colors*).

### The periplasmic pathway is closed in the presence of sodium

To establish which structure of BetP best represents the spectroscopic data measured in proteoliposomes, we defined a metric of the consistency of the simulated conformational ensemble with the PELDOR distributions. Specifically, we computed the work performed by EBMetaD to bias each simulation to reproduce the target distribution (**Figure 6**). We found that, for the data measured in the presence of sodium alone, simulations of the sodium-bound inward-facing structure required less work than simulations of the sodium-bound outward-facing structure to create an ensemble matching the experimental distribution (**Figure 6**).

**Figure 6.**
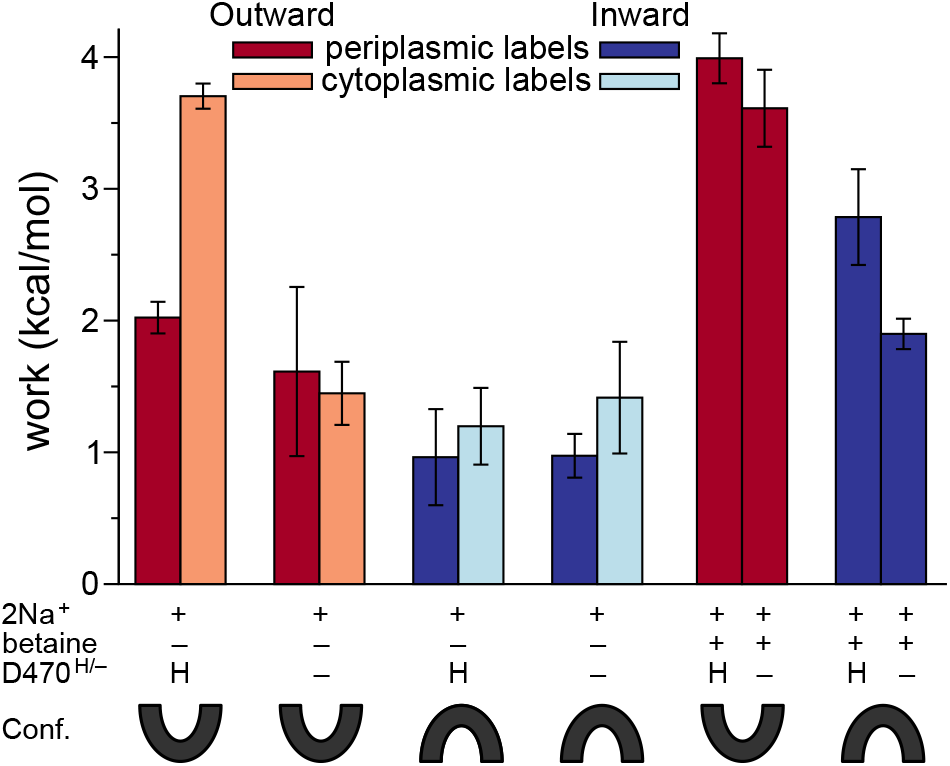
Work required by each simulation system to reproduce the experimental distance distribution. The work was computed by averaging the bias potential applied over the last 0.8 μs of each simulation of G450R5/S516R5 (periplasmic labels; *dark bars*), and S140R5/K489R5 (cytoplasmic labels; *light bars*). The simulations were carried out with two bound sodium ions (2Na^+^), with or without betaine, and with Asp470 either protonated (H), or deprotonated (−). The BetP conformation used to start the simulation was either inward- (PDB code: 4C7R, *blue-shaded bars*) or outward-facing (PDB code: 4LLH chain A, *red-shaded bars*). Error bars represent the standard deviation of work values obtained by averaging over the two halves of the last 0.8 μs.

Since G450 and S516 are on the periplasmic surface of the protein, the above result suggests that, in the presence of sodium, the periplasmic pathway is closed. However, given their location, these probes might equally be reporting on the closed state rather than the inward-open conformation simulated here. We therefore repeated the simulations using BetP monomers with probes attached on the cytoplasmic side at S140 and K489, whose distances were biased to the distribution obtained for S140R5/K489R5 in the presence of sodium (**Figure 3B, black line**). In this case, the protonated, outward-facing conformation required the largest magnitude of work (**Figure 4, light bars**). By contrast, simulations of the outward-facing state with Asp470 deprotonated, and both protonation states of the inward-facing state required less work to match the PELDOR distribution. However, it was not possible to distinguish between these three simulations within the error of the calculation. Based on this data, it would appear that in the presence of sodium alone the periplasmic probes are reporting on a closed periplasmic pathway, while the cytoplasmic probes are unable to distinguish whether the cytoplasmic pathway is open or closed. Since the closed state should not be populated in high sodium concentrations, we conclude that the preferred state of BetP under these conditions is the inward-facing conformation.

### In the presence of sodium and betaine, inward and outward conformations coexist

We next asked whether BetP has a preferred conformation under saturating conditions of both sodium and betaine, and how betaine alters that preference compared to sodium alone. In the presence of 300 mM NaCl and 5 mM betaine, the measured distance distribution for G450R5/S516R5 reveals a peak at distances of ~25 Å (**Figure 3D**). Molecular dynamics simulations of BetP bound to both betaine and sodium required more work to bias to the experimental distribution than was required for the equivalent systems with only sodium ions bound (**Figure 6**). This result suggests that no single starting structure accurately reflects the experimental ensemble obtained in the presence of betaine, consistent with the expectation that BetP can stochastically interconvert between inward- and outward-facing conformations when all required substrates are bound.

### Inward-facing state is the most compatible with sodium-only condition

To understand how the EBMetaD bias modifies the conformational preference for the labelled protein during the simulations, we de-biased the spin-label distance distribution obtained for the periplasmic probes (see Methods). The de-biasing procedure results in a distribution that mimics, approximately, the distances that might be obtained from a sufficiently long unbiased simulation, i.e., carried out with conventional molecular dynamics. Compared to the experimental distribution measured in the presence of sodium, which is >20 Å wide, and contains a large peak at ~43 Å, the distributions obtained after de-biasing are significantly sharper (**Figure 7A**). For the simulations of the outward-facing state, these peaks are centered around 32-37Å (**Figure 7A, red**), and therefore these trajectories poorly mimic the experimental distribution. This explains why additional bias was required to sample conformations in which the spin labels are further apart. In contrast, for the simulations of the inward-facing conformation, the spin label distance distributions obtained after de-biasing illustrate a better agreement with the width of the experimental distribution (**Figure 7A, blue**). Moreover, the distribution obtained for the simulation in which Asp470 was deprotonated illustrates that this system freely accesses conformations with a distance of ~43 Å, consistent with the experimental observation. The close correspondence of the experimental and debiased distributions may explain why the simulations of the inward-facing conformations required less work to achieve the experimental distance distribution (**Figure 6**).

**Figure 7.**
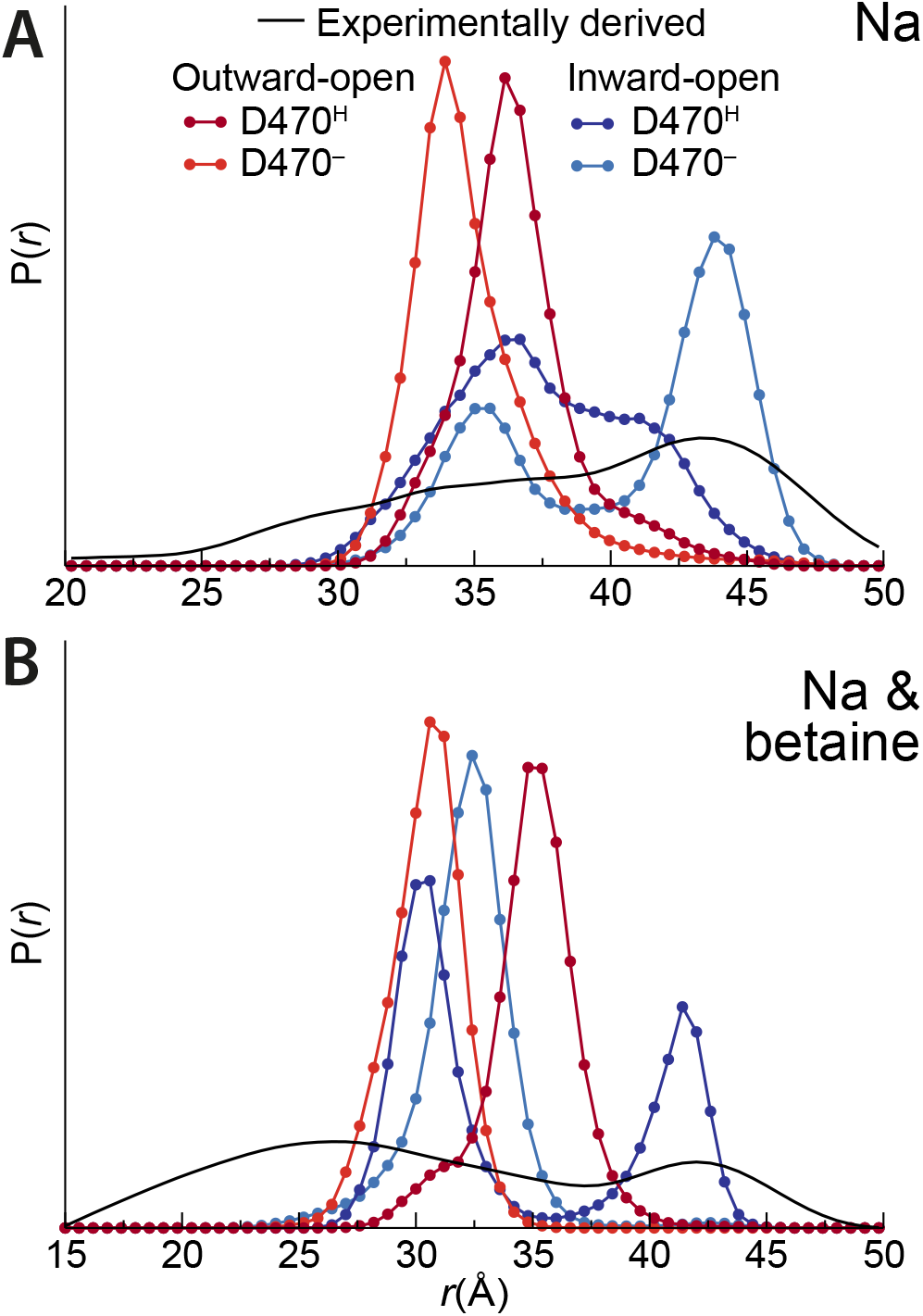
De-biased distance distributions of the periplasmic spin label pair. Simulations were carried out with either two sodium ions bound (**A**) or with two sodium ions plus the betaine substrate bound (**B**), with the protein in either an inward- (PDB code: 4C7R, *blues*) or outward-facing conformation (PDB code: 4LLH chain A, *reds*), and with Asp470 either protonated (*darker colors*), or deprotonated (*lighter colors*). The experimental distance distribution is shown for comparison (*black lines*), measured in the presence of either 500 mM NaCl (A) or 300 mM NaCl plus 5 mM betaine (B).

### No simulation encompasses the distribution obtained in the presence of betaine

In the presence of both betaine and sodium, the experimental distribution contains two distinct peaks and spans an even larger range of spin label distances (>35 Å) (**Figure 7B**). None of the simulations adequately matches either the range or the peak distribution for this data, which is reflected in the higher work needed to converge the experimental distance distribution under this condition, relative to the measurements performed in the presence of sodium alone. This suggests that the experimental distribution may reflect a mixture of states, or perhaps an additional state, such as the close state, not analyzed here.

### The bias acts primarily on the label orientation, not the protein backbone

Clearly, a spin-label distance distribution is not a simple measure of the sampled protein conformation, as it also reflects the relative dynamics of the spin labels. To examine whether the EBMetaD bias is primarily affecting the spin-label orientations or the protein backbone, we computed the angle between the spin labels, as well as the distance between the Cα atoms to which the labels are connected. In particular, we compare the results for these metrics obtained before and after de-biasing the trajectories. The Cα-Cα distance distributions are very similar in the biased and debiased trajectories, for all simulations, and under both conditions, indicating that EBMetaD was not strongly biasing the backbones of TM 8’ and 10’ (**Figure 8A-B**). Notably, these data illustrate how even for a single state of the transporter, the distance between these two helices fluctuates by 6-10 Å, which is larger than the difference between the equivalent distances in the structures (~2 Å).

**Figure 8.**
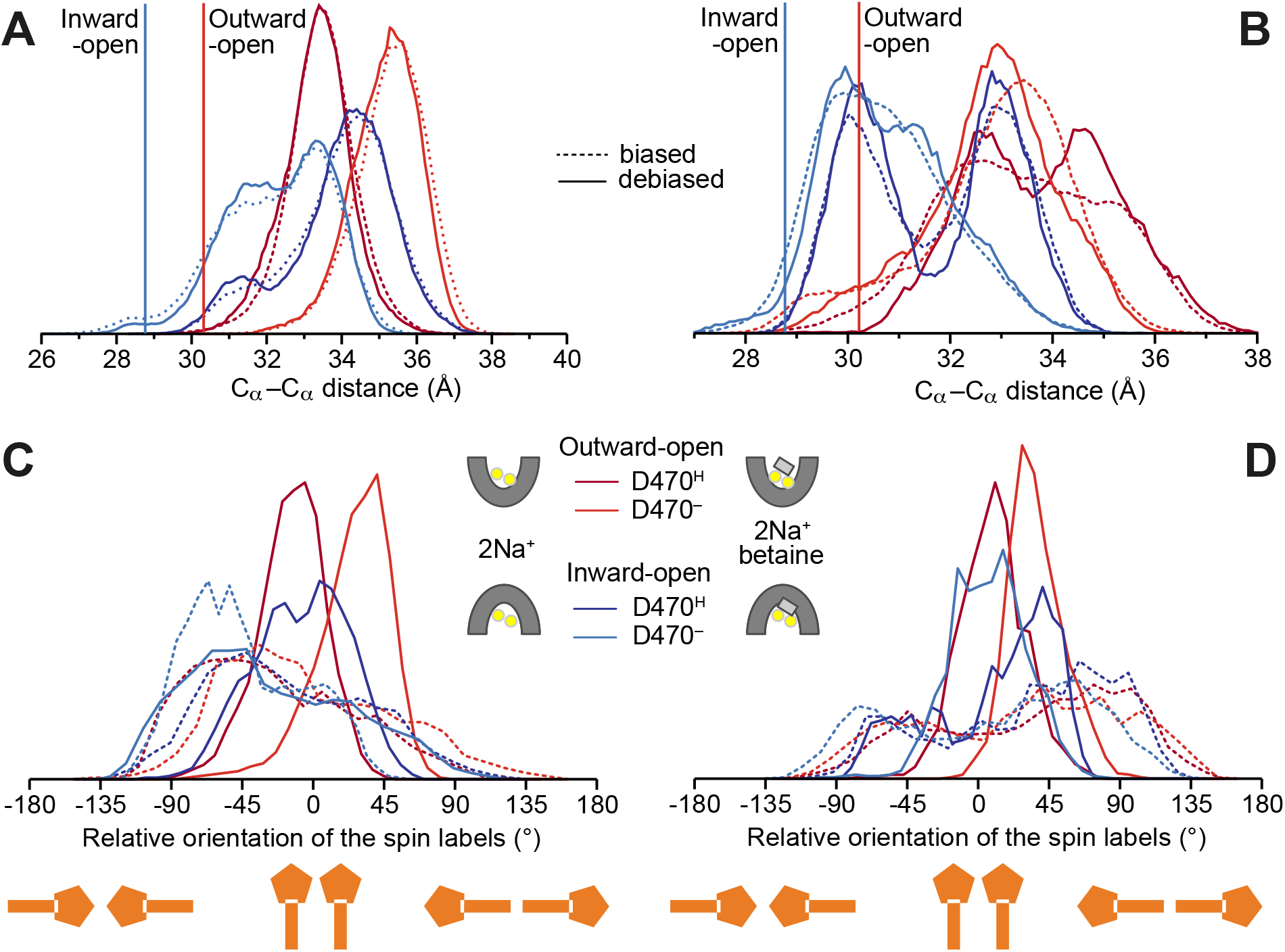
Analysis of backbone distances and spin label orientations. Distributions of spin label Cα-Cα distances and side chain orientations were calculated for the biased trajectories (*dashed lines*) and after de-biasing those same trajectories (*solid lines*) in simulations of G450R5/S516R5. (**A, B**) Distance between Cα atoms of the spin-labeled residues G450C and S516C during simulations of BetP with either two sodium bound (A) or two sodium and one betaine molecule bound (B). Vertical lines indicate the distances in the starting crystal structures. (**C, D**) Relative orientation of the spin labels during the simulations with two sodium bound (C) or with two sodium plus one betaine molecule bound (D). Data were obtained for simulations of BetP in outward- (PDB code: 4LLH chain A, *reds*), or inward-open conformations (PDB code: 4C7R, *blues*), with Asp470 either protonated (*darker colors*), or deprotonated (*paler colors*). The angle is computed for the axis connecting the backbone Cα and nitroxide N atoms in each spin label.

Unlike the Cα-Cα distances, the distribution of the angles between the spin labels is significantly different after de-biasing, for almost all of the simulated systems (**Figure 8C-D**). The exception is the simulation of the inward-open state with Asp470 deprotonated and in the presence of sodium alone (**Figure 8C**). Thus, in the background of 6-10 Å fluctuations of the protein backbone, sampling of the complete experimental distance distribution was achieved primarily by modifying the relative orientation of the spin labels.

## Discussion

The increasing availability of a range of structural and biophysical information for proteins such as BetP has revolutionized the membrane transport field, but has also presented challenges, in that different sources of data often appear to be incompatible or not clearly consistent. In this context, novel strategies to evaluate and integrate different kinds of experimental data with atomistic, dynamic representations of these proteins and their environments have the potential to drive unprecedented mechanistic insights. In the case of BetP, for example, the small distance changes implied by comparison of the outward- and inward-open crystal structures cannot be readily reconciled with the broad distance distributions obtained for spin labels attached to the protein. However, fully atomistic molecular simulations of the spin-labelled protein, based on the enhanced-sampling technique known as EBMetaD, enabled us to assess which of the existing structures is most representative of the molecular ensembles captured in the experimental distance distributions. The results show that the distributions measured in the presence of sodium are most compatible with conformations in which the periplasmic pathway is closed, and thus, most likely, the inward-open states. This result is non-trivial in that the physiological role of BetP is to capture betaine from the extracellular solution, which would suggest a preference for the outward-facing state in the presence of sodium, like LeuT (18, 19). Nevertheless, our observations for BetP are consistent with EPR measurements on Mhp1 (14, 17), and with the fact that BetP crystallizes preferentially in inward-facing conformations in saturating sodium concentrations (16).

### Sensitivity of the methodology to the experimental distribution

In this study, the EBMetaD methodology was used to reproduce distributions that had been obtained by Tikhonov regularization of the PELDOR time traces. However, the Tikhonov regularization does not result in a unique solution and therefore it is possible that the magnitude of the biasing work applied during the simulations would differ if a different solution were to be considered. A similar concern relates to the postprocessing of the periplasmic spin-label distributions to remove inter-protomer contributions. To assess the extent to which our conclusions might be dependent on these choices, we recomputed the biasing work that is required to enforce the experimental distributions on the periplasmic pair, but only for distances <37 Å, i.e., excluding the contributions from the peak at ~42 Å that is present in both conditions (**Figure 3D**). This analysis reproduced the trends observed for the entire distribution (**Figure 9**). First, simulations initiated with the inward-open structure required less work to reproduce the target distributions, both in the presence and absence of betaine. And second, the simulations in the presence of both sodium and betaine required more work to reproduce the corresponding distribution than the simulations in the presence of sodium alone. Thus, although the specific value of the work applied may vary somewhat, the overall strategy appears not to be sensitive to the features of the peak at ~42 Å; thus, it is also unlikely to be sensitive to variations in the solution of the Tikhonov regularization procedure.

**Figure 9.**
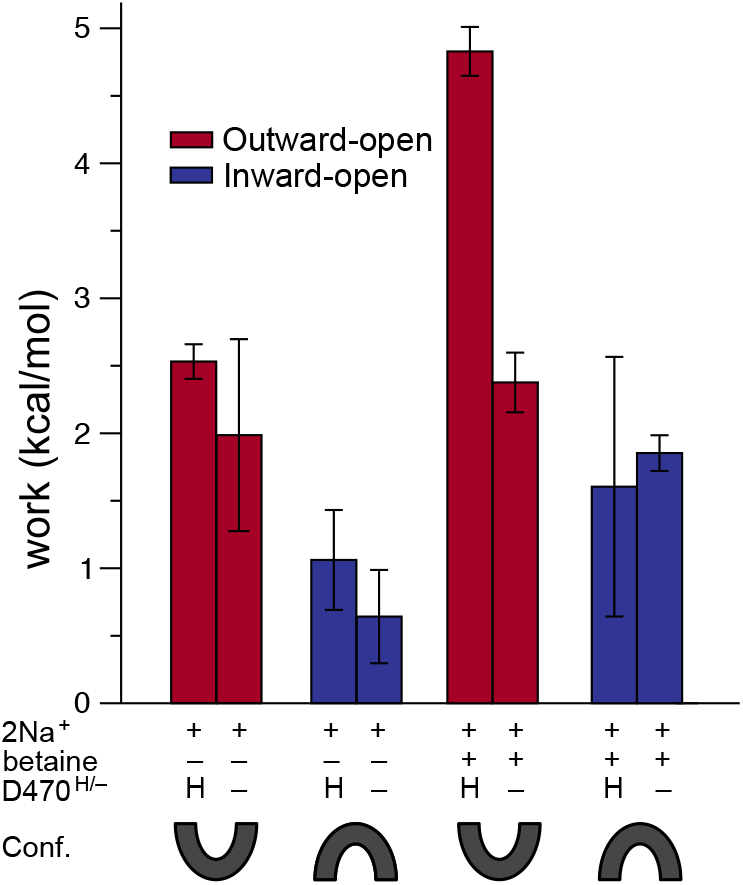
Work required by each BetP simulation system to reproduce a truncated experiment distance distribution. The work was computed by averaging the bias potential applied for distances <37 Å (see Methods), during the last 0.8 μs of each simulation of G450R5/S516R5 (periplasmic labels), with two bound sodium ions (2Na^+^), with or without betaine, and with Asp470 either protonated (H), or deprotonated (−). The initial protein conformation was either inward- (PDB code: 4C7R, *blue bars*) or outward-facing (PDB code: 4LLH chain A, *red bars*). Error bars represent the standard deviation of work values obtained by averaging over the two halves of the last 0.8 μs.

### The role of Asp470 in transport

The protonation state of Asp470 computed using electrostatics calculations was sensitive to the conformation of the protein and the presence or absence of its substrates, suggesting that the transport cycle of BetP entails protonation and deprotonation of this residue. Since proton gradients are unable to drive betaine uptake for BetP, however, we propose that Asp470 binds and releases protons from the same side of the membrane, namely the cytoplasm. Specifically, our calculations suggest that release of betaine and sodium from the inward-facing state is fostered by protonation of Asp470, and that this proton has to be released back into the cytoplasm for BetP to reset the transport cycle via the closed and outward-open apo states. It is worth noting that Asp470 is conserved among sodium-dependent transporters of the BCCT family (6) and therefore may play a role in sodium recognition. It will be interesting to explore the pH dependence of transport in BetP and its relationship with the nature of the side chain at position 470.

It could be argued that the conformational preference of BetP in crystals reflects the low pH (5.5) used in the crystallization conditions, which would favor protonation of Asp470 and thereby inward-facing states. However, the results from the simulations show that the transporter adopts an inward-facing conformation in the presence of sodium even at pH 7.5, which is the condition of the PELDOR measurements. Therefore, low pH may instead be required to stabilize crystal contacts involving other ionizable residues in the protein.

### The role of concentration gradients

The PELDOR measurements carried out here utilized equal concentrations of substrates on either side of the proteoliposome membrane. The reason for this choice is that the orientation of BetP is unknown, and therefore this protocol ensures that all proteins will experience the same environment. As mentioned, the observation that BetP prefers outward-closed (or inward-open) conformations in the presence of sodium is not unprecedented, and is in line with earlier results for Mhp1. Nevertheless, we cannot rule out the possibility that BetP and Mhp1 would instead prefer outward-open states if an inward-oriented sodium gradient were applied. Assuming that a gradient would not alter the conformational preference observed in proteoliposomes, the question that arises is why BetP is not primarily outward-facing in the presence of sodium. An intriguing possible explanation relates to the activation of transport in response to osmotic stress. This activation process is triggered by an increase in the intracellular potassium concentration, and accelerates transport by up to two orders of magnitude (25, 43). If the rate-limiting step of the transport cycle is the resetting of apo BetP from the inward-to the outward-facing state, then activation might be achieved simply by changing the relative stabilities of these two states, and/or by reducing the kinetic barrier to the closure of the intracellular pathway. Measurements of the effect of potassium on the conformational equilibrium may help examine this possibility. However, if measured differences in the presence of potassium fall within the range of the probe flexibility, it will not be possible at this stage to use molecular simulations to help interpret those data, because the mechanism of activation involves intratrimer interactions between cytoplasmic segments whose structures remain poorly characterized.

In summary, this study provides insights into transport cycle of BetP and how the conformational preferences of this transporter are altered upon recognition of its substrates. More broadly, this work illustrates the potential of EBMetaD simulations to facilitate an integrative, molecular-level interpretation of structural and spectroscopic data on membrane proteins.

## Supplementary Figures

**Supplementary Figure 1:**
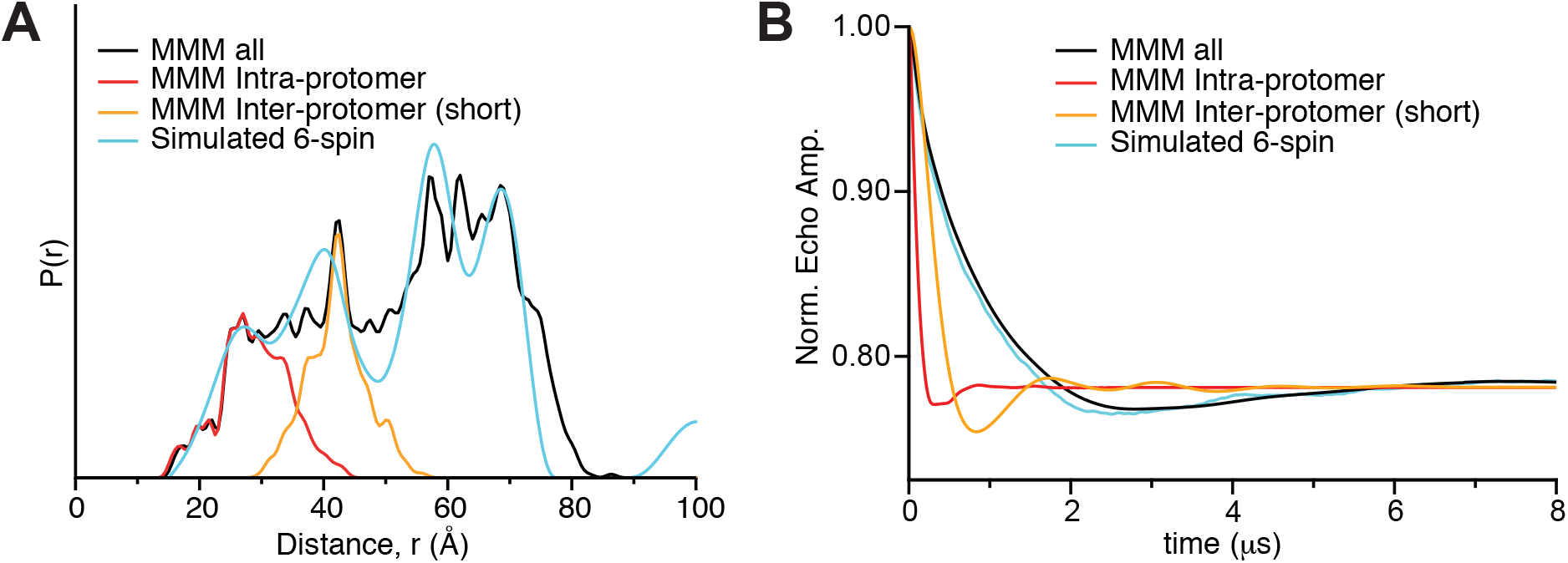
Simulated PELDOR data examining the likelihood of ghost peaks due to multi-spin effects. (**A**) Distance distribution extracted from predicted conformational ensembles of BetP spin labels at positions 450 and 516 on the trimer (PDB 4DOJ). The distances were estimated using MMM with the ‘R1A_298K_xray’ rotamer library. We then extracted two-spin distances for all predicted 2-spin distances (*black*), for all pair-wise distances within a protomer (*red*), and for the shortest distribution of pair-wise distances between two probes in different protomers (*orange*). The simulated 6-spin distance distribution (*cyan*) was reverse-engineered from the predicted PELDOR time trace shown in (**B**) using DeerAnalysis with ‘ghost spin’ scaling of 6 spins. The Tikhonov regularization was carried out with a smoothing parameter of 1000. (**B**) PELDOR time traces predicted from the MMM spin label distributions shown in (**A**), assuming either six spins (*cyan*), or only two-spin contributions (*black, red* and *orange*).

**Supplementary Figure 2.**
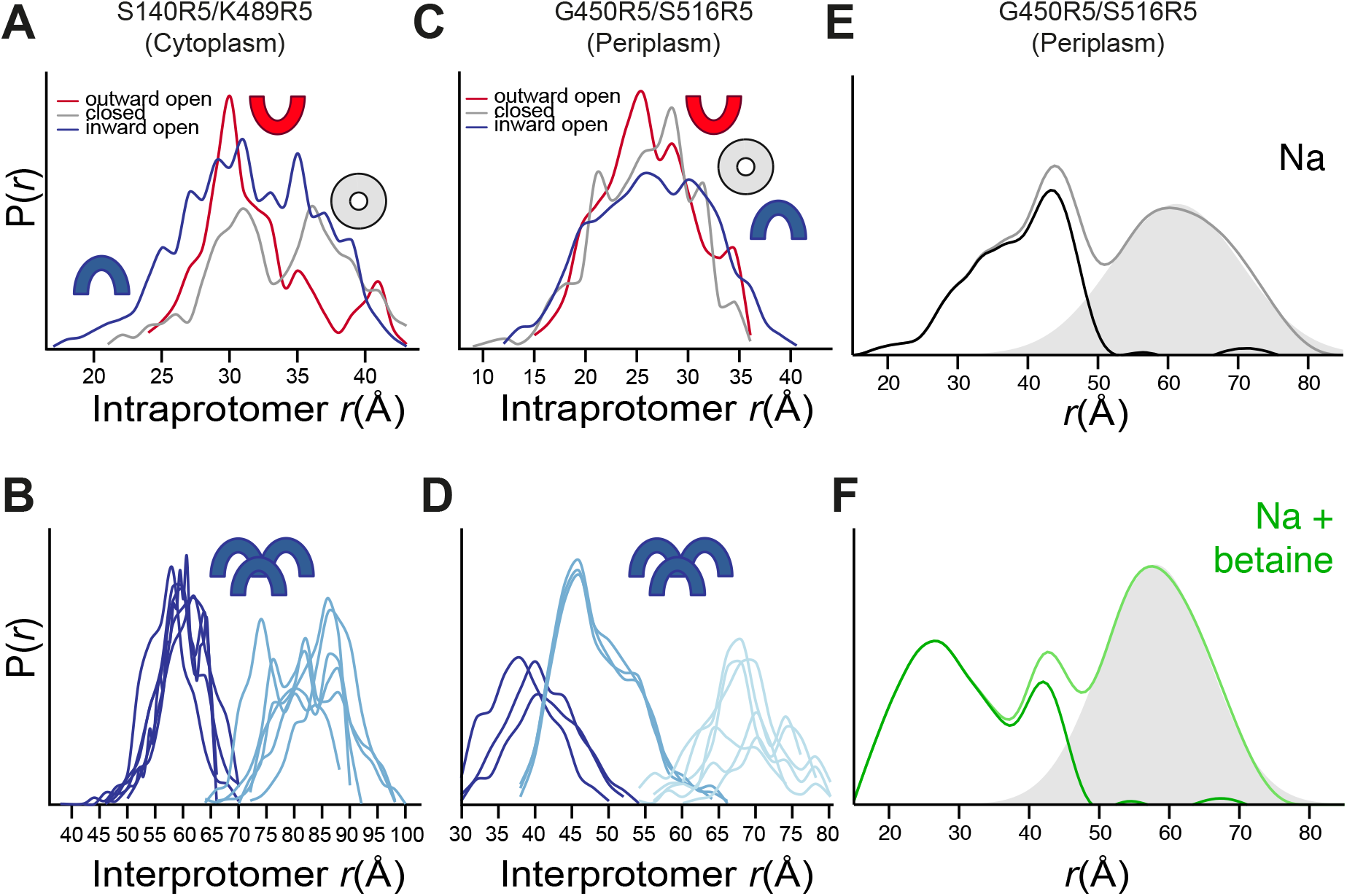
Spin label distance distributions predicted by sampling spin label rotamers on static crystal structures, and filtering of long-range peaks. Spin label rotamer-library based predictions were conducted using the MMM package (21, 23) for individual protomers in one of three conformational states (**A, C**), or a trimer in the inward-facing state (**B, D**). Probes were attached at cytoplasmic positions S140 and K489 (A, B), or periplasmic positions G450 and S516 (C, D). Lines represent the distances obtained for all rotamers of a single pair of labels. In (B) and (D), different shades of blue highlight distinct populations observed at different distance ranges. (**E, F**) Filtering out of intra-protomeric distances from the periplasmic spin label distance distributions measured in the presence of either 500 mM NaCl (E), or 300 mM NaCl and 5 mM betaine (F). Based on the spin-label rotamer predictions (A-D), the peaks observed at ~60 Å must represent coupling between probes on adjacent protomers. Therefore, we subtracted this peak from the PELDOR-derived distance distribution for use with simulations of the BetP monomer. Specifically, a Gaussian function centered at this peak (*shaded surface*) was subtracted from the experimental data (*darker lines*) to obtain the final distribution (*lighter lines*).

## Supporting Information

### Processing the PELDOR data for multi-spin contributions

In the configuration of two spin labels on each protomer, BetP contains six spin labels per trimer. Aside from the large number (i.e., 15) of expected different spin-spin interactions, the presence of multiple labels also raises the possibility of more than two spins coupling with each other. That is, a three (or more) spin system could not only lead to the expected 3 distances, but also to additional ghost distances due to trigonometric function with non-linear relation. These ghost distances would be due to larger (artificial) frequency and therefore the ghost peaks would show up at shorter distances than expected. The probability of observing such ghost distances depends on the modulation depth of the signal, which in turn depends on the power of the applied microwave. While more ‘modern’ Q-Band spectrometers with high 150W microwave power could result in depths of 50% or more, we obtained data with relatively low modulation depth values (~20%), which helped to reduce multi-spin signals. The second technique used to reduce such ghost peaks is power scaling of the recorded PELDOR time traces (1). Specifically, the normalized time trace is scaled by the power of 1/(*n*-1), where the contributing number of spins is n. In practice, this was accomplished by analyzing the PELDOR time traces using the ‘ghost spin’ option in the DeerAnalysis software, rescaling them by the power of 1/5, assuming six spins.

### Assessing the contribution of multi-spin dipolar couplings

To assess the likely contribution from multi-spin dipolar couplings, we used MMM 2013 to model the rotamer states of the MTSSL spin label on positions 450 and 516 of an X-ray structure of the BetP trimer (PDB entry 4DOJ). First, to identify the contributions from intra- and inter-protomer spin label pairs that would occur in the absence of multi-spin coupling, we modeled specific pairs of spin labels and computed the corresponding spin-spin distances, as well as the predicted PELDOR time traces for various combinations thereof. Second, we generated models of the BetP trimer containing all six spin labels, for which we computed the corresponding distance distributions and the PELDOR time traces, including all multiple spin effects that are theoretically possible. This predicted PELDOR trace should be directly comparable with the experimental data. To identify the ghost peaks in the MMM 6-spin distance distribution, the predicted PELDOR time traces were then analyzed with DeerAnalysis to reverse engineer a distance distribution, using the same power-scaled approach used for the experimental data. Specifically, the simulation assumed a modulation depth of 22% and, prior to applying Tikhonov regularization, the time traces were rescaled to the power of 1/5, to reduce ghost effects.

The MMM-simulated PELDOR distance distributions and time traces are shown in **Figure S1**. Importantly, the complete distance distribution of the MMM 6-spin structural ensemble (*black*, **Figure S1A**) is very similar to the reverse-engineered distribution from a simulated PELDOR time trace in which all multiple spin effects were taken into account (*cyan*, **Figure S1A**). In particular, no additional peak in the distance distribution at short distances, i.e., shorter than the intra-protomer distances (*red*, **Figure S1A**), was generated when incorporating the multi-spin effect. We note that although some variation is visible at larger distances, such effects would be negligible for time traces containing noise at levels encountered in typical experiments.

### Assigning the protonation state of ionizable residues in BetP

Classical molecular dynamics simulations do not allow for bond breaking or formation, including protonation or deprotonation of ionizable side chains. Therefore, the ionization states of such residues must be assigned at the start of the simulations. Continuum electrostatics calculations previously revealed two ionizable residues buried in the core transmembrane region of the protein whose protonation states might be non-standard at neutral pH, namely Glu161 from TM 1’ and Asp470, from TM 8’ (2). However, these calculations were carried out only using the outward-facing structure. Since the pKa of a given residue can depend on the conformation of the protein around it, we therefore extended the earlier calculations to consider also the other major states of the transport cycle that have been characterized crystallographically (**Figure 4, Table S2**).

The pKa calculations clearly indicated that Glu161 is likely to be protonated at pH 7 in all conformational states of BetP, with the only exception being the substrate- and sodium-bound outward-facing conformation when assuming a dielectric constant for the protein, ε of 8 (**Table S2**). This result might reflect the proximity of Glu161 to Gly153, which was mutated to aspartate in this crystal structure (2) and whose charge could modify the conformation or hydration close to Glu161. Since we simulate the Gly153 (WT) construct, we assign Glu161 as charged in all simulations.

Unlike the results obtained for Glu161, the pKa of Asp470 appears to be sensitive to the conformation of the protein. Specifically, the side-chain of Asp470 is predicted to be charged in the outward-facing and closed states (**Figure 4B, Table S2**), in which the cytoplasmic pathway is closed, and the aspartate sidechain is oriented towards Ser471 and the Na2 site (**Figure 4A**). In contrast, in the inward-facing states (**Figure 4B, Table S2**), when the sidechain is oriented toward Ser293 in the TM4’-5’ loop (**Figure 4A**), the pKa of Asp470 shifts to values of ~7. The computed pKa values of Asp470 indicate that its protonation state could change during the transport cycle and, importantly, it could differ between the inward- and outward-facing conformations studied here.

In summary, we assume Glu161 is protonated, but we cannot assign a single protonation state for Asp470.

**Supplementary Table 1:**
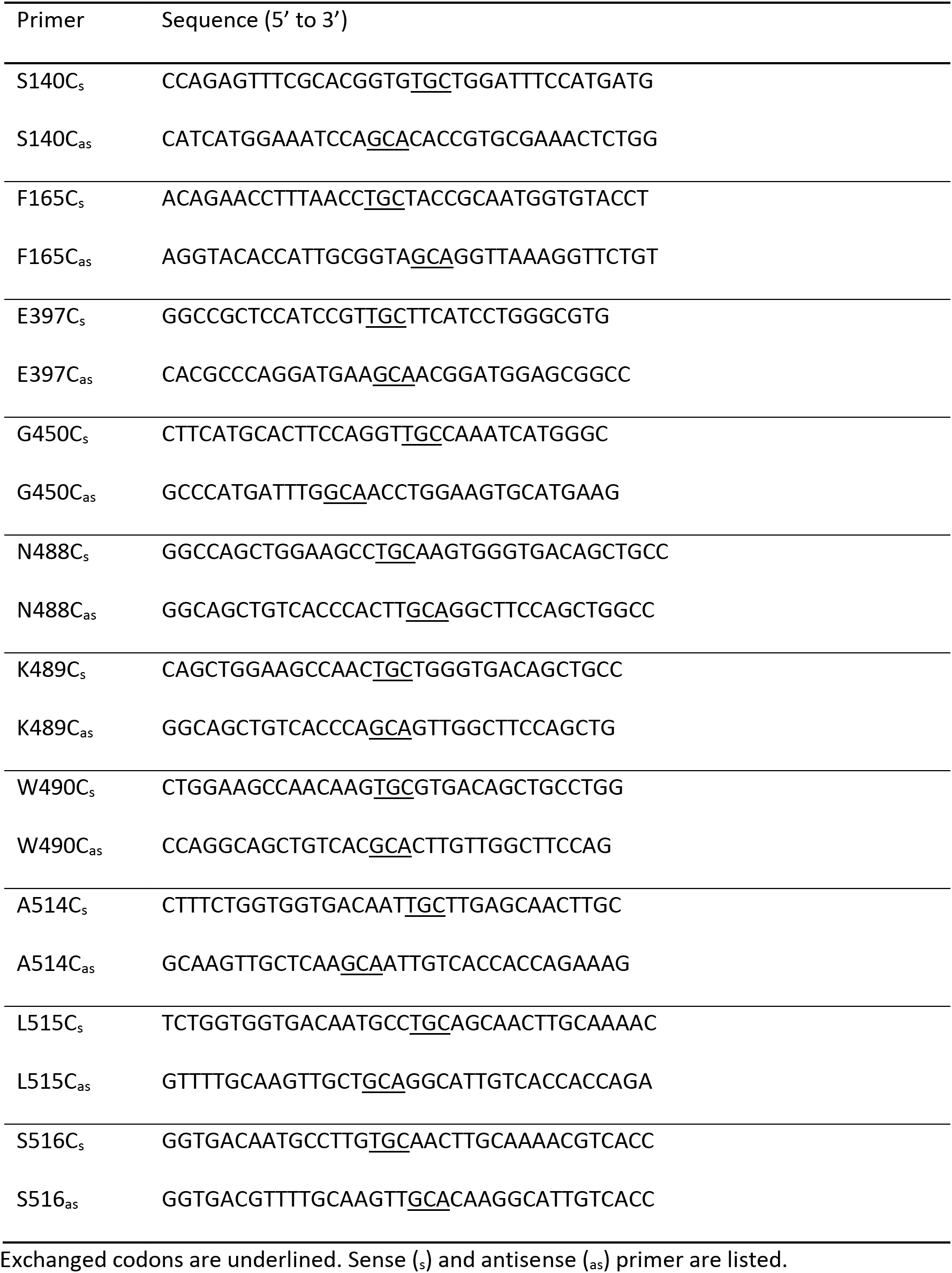
Primers.

**Supplementary Table 2:** Computed pKa values for ionizable residues in the transmembrane region of BetP.

| Conformation | PDB | Residue |  |  |  |
| --- | --- | --- | --- | --- | --- |
|  |  | E161 |  | D470 |  |
| | | $\epsilon=4$ | $\epsilon=8$ | $\epsilon=4$ | $\epsilon=8$ |
| **Outward + betaine + 2Na<sup>+</sup>** | **4LLH_B\*** | **8** | **6** | **4** | **3** |
| Closed + betaine + 2Na<sup>+</sup> | 4AIN_B | 10 | 7 | 3 | 3 |
| **Inward + betaine + 2Na<sup>+</sup>** | **4AIN_C** | **10** | **8** | **7** | **6** |
| **Inward** | **4C7R_C** | **11** | **8** | **8** | **6** |
| Closed | 4AIN_A | >14 | 9 | 6 | 5 |
| **Outward** | **4LLH_B** | **10** | **8** | **7** | **6** |

pKa values were computed for different structures in order of the steps in the physiological transport cycle, with the continuum electrostatics method MCCE (3) and with the protein dielectric constant set to 4 or 8. In bold are highlighted the results for the inward- and outward-facing conformations, as distinct from the closed states, in which both pathways are occluded.
\*Data for the outward-facing state are taken from (2), and were obtained from the crystal structure of G153D mutant that was *in silico* mutated back to the wildtype sequence, while the choline molecule was substituted by a betaine molecule.

**Supplementary Table 3:**
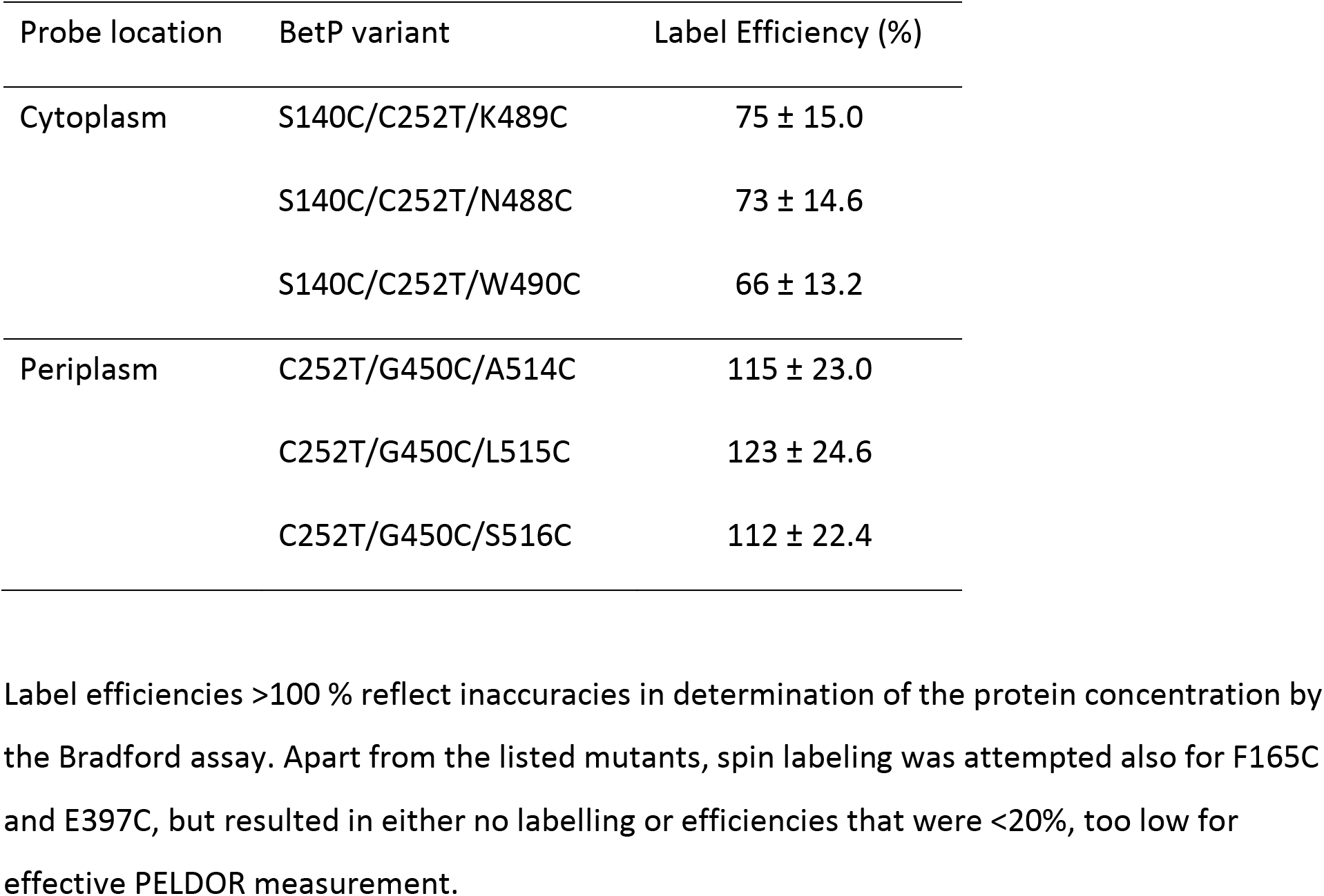
Efficiency of spin labeling of BetP variants.

| Probe location | BetP variant | Label Efficiency (%) |
| --- | --- | --- |
| Cytoplasm | S140C/C252T/K489C | 75 ± 15.0 |
|  | S140C/C252T/N488C | 73 ± 14.6 |
|  | S140C/C252T/W490C | 66 ± 13.2 |
| Periplasm | C252T/G450C/A514C | 115 ± 23.0 |
|  | C252T/G450C/L515C | 123 ± 24.6 |
|  | C252T/G450C/S516C | 112 ± 22.4 |

Label efficiencies >100 % reflect inaccuracies in determination of the protein concentration by the Bradford assay. Apart from the listed mutants, spin labeling was attempted also for F165C and E397C, but resulted in either no labelling or efficiencies that were <20%, too low for effective PELDOR measurement.

## Footnotes

1 Because BetP contains two additional helices before the conserved core, we use helix numbering (−2, −1, 1 −10) corresponding to LeuT for ease of comparison.

## Acknowledgements

This research was supported in part by the Division of Intramural Research of the NIH, National Institute of Neurological Disorders and Stroke, the Deutsche Forschungsgemeinschaft Sonderforschungsbereich SFB 807, and the Max Planck Institute of Biophysics, and utilized the computational resources of the NIH HPC Biowulf cluster (http://hpc.nih.gov) and the LOBOS network at the National Heart, Lung and Blood Institute. We thank José Faraldo-Gómez and Fabrizio Marinelli for useful discussions.

